## Supplementary Data for "Testing the adaptive walk model of gene evolution"

**Tables**

**Table S1.** Kendall's correlation coefficients for the relationship between the mean value of each co-factor in each age class and ω, $\omega_{na}$ and $\omega_a$ for each "high" and "low" group. The Kendall's correlation coefficients for the relationship between the co-factor and gene age in each "high" and "low" group are presented in the column "co-factor~Age".

| | *Co-factor* | *Category* | $\omega$ | $\omega_{na}$ | $\omega_a$ | *co-factor ~ Age* |
|---|---|---|---|---|---|---|
| 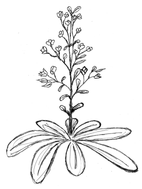 | **RSA** | High | 0.6667 * | 0.6111 * | 0.6667 * | 0.5434 ** |
| | | Low | 0.5000 (.) | 0.5000 (.) | 0.2778 | 0.6381 *** |
| | **Disorder** | High | 0.7576 *** | 0.8181 *** | 0.3939 (.) | 0.5049 ** |
| | | Low | 0.1515 | 0.1515 | -0.1212 | 0.2000 |
| | **Length** | Long | -0.7333* | -0.6000 (.) | 0.2000 | -0.7903 *** |
| | | Short | -1.0000 ** | -0.8667 * | -0.6000 (.) | -0.394 * |
| | **Expression** | High | 0.0667 | 0.0667 | 0.2000 | -0.096 |
| | | Low | -0.8667 * | -1.0000 ** | -0.3333 | -0.842 *** |
| 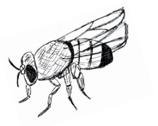 | **RSA** | High | 0.5272 * | 0.7091 ** | 0.4545 (.) | 0.757 *** |
| | | Low | 0.2363 | 0.0909 | 0.3818 | 0.484 * |
| | **Disorder** | High | 0.3636 (.) | 0.3939 (.) | 0.2424 | 0.454 * |
| | | Low | -0.1515 | -0.1515 | -0.0303 | -0.424 (.) |
| | **Length** | Long | -0.0667 | -0.3333 | -0.0667 | -0.363 (.) |
| | | Short | -0.7333 * | -0.8667 * | -0.4667 | -0.455 * |
| | **Expression** | High | -0.2000 | -0.0667 | -0.4667 | -0.259 |
| | | Low | -0.6000 (.) | -0.7333 * | -0.7333 * | -0.419 (.) |

**Note.** For each variable, the correlation coefficients are shown with the respective significance (*$P < 0.05$; **$P < 0.01$; ***$P < 0.001$; "." $0.05 \leq P < 0.10$) in *Arabidopsis* and *Drosophila*.

**Table S2.** Linear models' estimates when using the rates of protein evolution (ω, $\omega_{na}$ and $\omega_a$) as response variables and the mean value of the cofactor per category as a putative explanatory variable, together with gene age, the species (*Arabidopsis* or *Drosophila*), cofactor-category, and their interaction with gene age.

| Co-factor | Variable | $\omega$ | $\omega_{na}$ | $\omega_a$ |
|---|---|---|---|---|
| **Gene length** | (Intercept) | 0.137 (***) | 0.027 | 0.111 (***) |
| | **GeneAge** | **0.028 (***)** | **-0.001** | **0.029** (***) |
| | speciesDrosophila | -0.032 | -0.021 | -0.005 |
| | categoryshort | 0.197 | 0.264 (*) | -0.131 (**) |
| | **GeneAge:speciesDrosophila** | **-0.008 (*)** | **0.012 (**)** | **-0.022** (***) |
| | categorylong:mean.length | 0.000 | 0.000 | 0.000 (**) |
| | categoryshort:mean.length | 0.000 | 0.000 (*) | 0.000 (*) |
| | GeneAge:categoryshort | -0.010 (.) | -0.007 | |
| **Gene expression** | (Intercept) | 0.087 (***) | 0.046 (*) | 0.042 (.) |
| | **GeneAge** | **0.026** (***) | **0.007 (***)** | **0.019 (***)** |
| | speciesDrosophila | -0.025 | 0.046 (*) | -0.065 (**) |
| | categoryLow | 0.099 (**) | -0.024 | 0.164 (**) |
| | categoryHigh:mean.exp | 0.000 | -0.001 (.) | 0.001 (.) |
| | categoryLow:mean.exp | -0.008 | -0.002 | -0.012 |
| | **GeneAge:speciesDrosophila** | **-0.012 (***)** | | **-0.010 (***)** |
| | GeneAge:categoryLow | | | -0.004 |
| **Protein disorder** | (Intercept) | 0.060 (.) | 0.431 (*) | -0.382 (*) |
| | **GeneAge** | **0.051 (***)** | **0.027 (***)** | **0.023 (***)** |
| | speciesDrosophila | 0.063 | 0.071 (.) | -0.001 |
| | categorylow | -0.047 (*) | 0.057 | -0.386 |
| | **GeneAge:speciesDrosophila** | **-0.034 (***)** | **-0.015 (*)** | **-0.018 (***)** |
| | categoryhigh:mean.dis | | -4.288 (*) | 4.370 (**) |
| | categorylow:mean.dis | | -7.299 | 11.349 (*) |
| | GeneAge:categorylow | | -0.010 | 0.011 (*) |
| **Relative solvent accessibility** | (Intercept) | 0.101 (***) | -0.006 | 0.259 (.) |
| | **GeneAge** | **0.036 (***)** | **0.012 (***)** | **0.026 (***)** |
| | speciesDrosophila | -0.030 | 0.035 (*) | -0.069 (**) |
| | **GeneAge:speciesDrosophila** | **-0.015 (**)** | | **-0.014 (***)** |
| | categoryLow | | | 0.367 |
| | categoryHigh:mean.rsa | | | -0.309 |
| | categoryLow:mean.rsa | | | -1.582 (*) |

**Note.** For each variable, the correlation coefficients are shown with the respective significance (*$P < 0.05$; **$P < 0.01$; ***$P < 0.001$; "." $0.05 \leq P < 0.10$).

**Table S3.** MK regression estimates, z-scores, and respective p-values.

| | *Parameter* | *Estimate* | *SE* | *Z-score* | *p-value* |
|---|---|---|---|---|---|
| 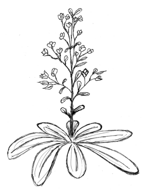 | Protein Length | -0.3646 | 0.0859 | -4.2470 | 2.1665e-05 |
| | Gene Expression | -0.2981 | 0.0333 | -8.9434 | 3.7748e-19 |
| | **Gene Age** | **0.2013** | **0.0184** | **10.9155** | **9.7231e-28** |
| | Protein Disorder | 18.6398 | 1.8520 | 10.0646 | 7.9178e-24 |
| | RSA | 0.9794 | 0.5328 | 1.8381 | 6.6047e-02 |
| | Grantham's distance | 1.6449 | 0.2349 | 7.0027 | 2.5116e-12 |
| | Protein Function | 10.9524 | 24.5421 | 0.4463 | 6.5540e-01 |
| 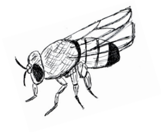 | Protein Length | -0.0404 | 0.0233 | -1.7348 | 8.2785e-02 |
| | Gene Expression | -0.0255 | 0.0150 | -1.6999 | 8.9147e-02 |
| | **Gene Age** | **0.0529** | **0.0097** | **5.4649** | **4.6311e-08** |
| | Protein Disorder | 4.3035 | 0.7033 | 6.1193 | 9.3989e-10 |
| | RSA | -0.0209 | 0.2600 | -0.0805 | 9.3582e-01 |
| | Grantham's distance | 4.0508 | 0.0976 | 41.4829 | 0.0000e+00 |
| | Protein Function | -1.7831 | 0.0867 | -20.5715 | 4.9403e-94 |
| | ChrX | -0.2530 | 0.0979 | -2.5850 | 9.7370e-03 |

**Table S4.** Partial $R^2$ estimates for each factor analysed in the MK regression analysis.

| | *Variables* | *Log likelihood (reduced)* | *Log likelihood (full)* | *Partial $R^2$* | *Relative partial R2(1)* |
|---|---|---|---|---|---|
| 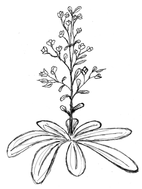 | Protein Function | -1730085.78830 | -1704071.01001 | 7.2821e-03 | 100% |
| | Grantham's distance | -1724125.91445 | -1704071.01001 | 5.6185e-03 | 77% |
| | Gene Expression | -1706525.87131 | -1704071.01001 | 6.8945e-04 | 9% |
| | Protein Length | -1704596.03241 | -1704071.01001 | 1.4749e-04 | 2% |
| | **Gene age** | **-1704165.50280** | **-1704071.01001** | **2.6547e-05** | **0.4%** |
| | Protein Disorder | -1704119.58878 | -1704071.01001 | 1.3648e-05 | 0.2% |
| | RSA | -1704084.06287 | -1704071.01001 | 3.6671e-06 | 0.05% |
| 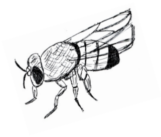 | Grantham's distance | -383465.345328 | -375682.00942 | 6.2597e-03 | 100% |
| | Protein Function | -379470.395391 | -375682.00942 | 3.0517e-03 | 49% |
| | Protein Length | -375960.543575 | -375682.00942 | 2.2469e-04 | 4% |
| | Gene Expression | -375870.540657 | -375682.00942 | 1.5209e-04 | 2% |
| | Protein Disorder | -375732.957146 | -375682.00942 | 4.1102e-05 | 0.7% |
| | Chr X | -375718.212571 | -375682.00942 | 2.9207e-05 | 0.5% |
| | **Gene age** | **-375707.013831** | **-375682.00942** | **2.0174e-05** | **0.3%** |
| | RSA | -375705.886390 | -375682.00942 | 1.9263e-05 | 0.3% |

(1) Partial R2 is relative to the partial R2 of the strongest effect. These values were estimated using 7,118,791 and 2,479,018 sites in *Arabidopsis* and *Drosophila*, respectively.

### Figures

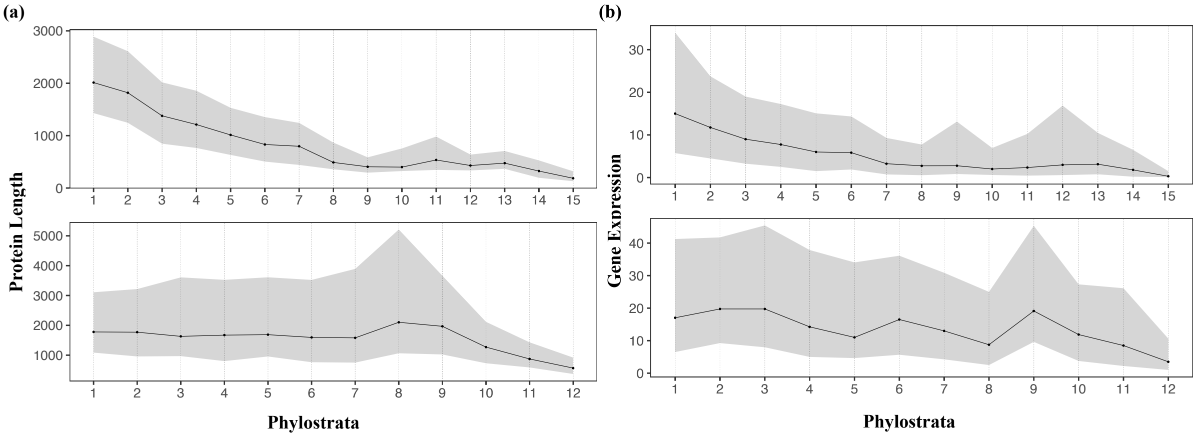


**Figure S1**. Relationship between gene age and gene length (a) and gene expression (b) for *A. thaliana* (top) and *D. melanogaster* (bottom). This analysis was performed by categorizing gene age according to the clades defined in Figure 1a. For each clade, the median value of gene length and gene expression is depicted with the black dot. The shaded area represents the values of gene length and mean expression levels within the 1st and 3rd quartile.

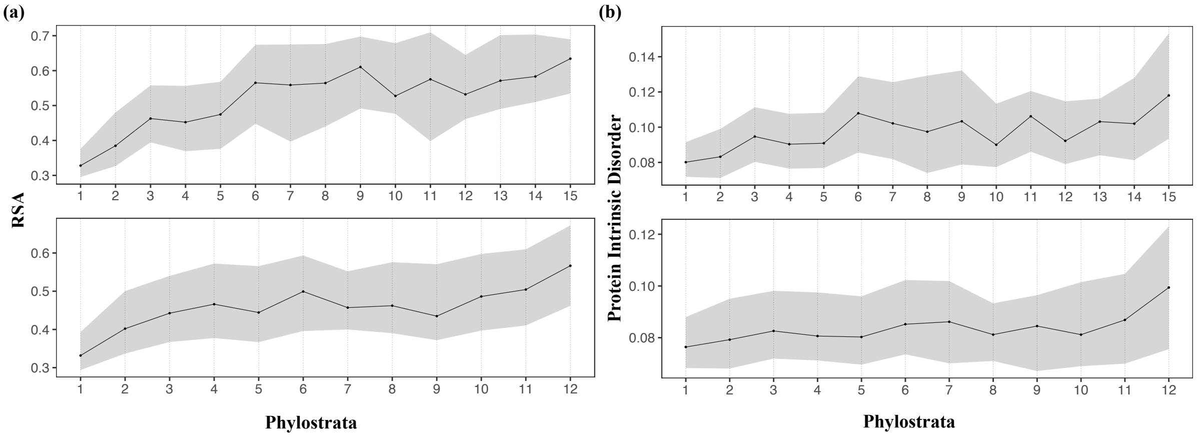


**Figure S2**. Relationship between gene age and RSA (a) and protein intrinsic disorder (b) for *A. thaliana* (top) and *D. melanogaster* (bottom). Legend as in Figure S1.

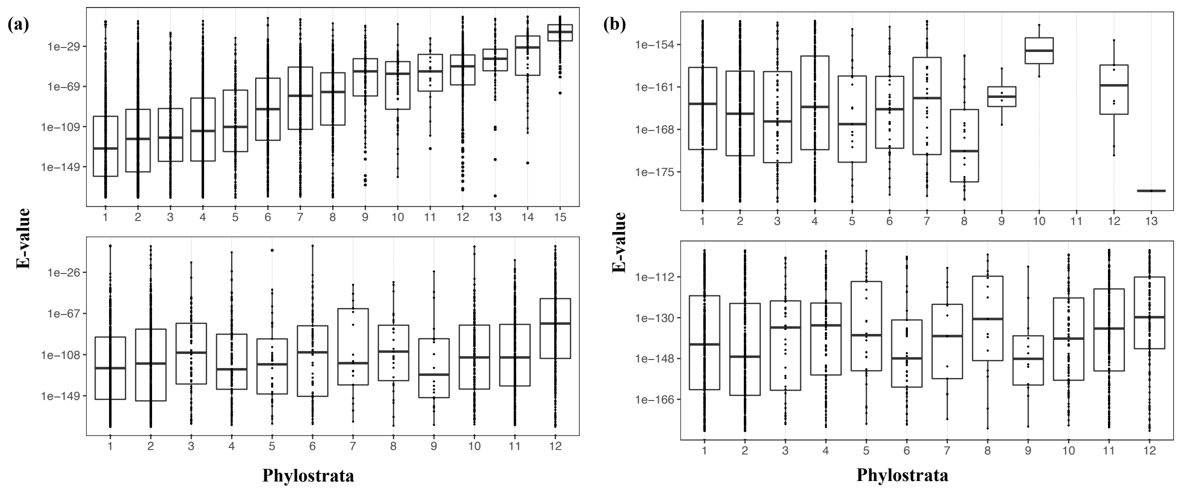


**Figure S3**. Relationship between gene age and E values before (a) and after (b) the E value correction for *A. thaliana* (top) and *D. melanogaster* (bottom). Each black dot represents a gene and median E value for each clade is represented with the black line in the boxplot.

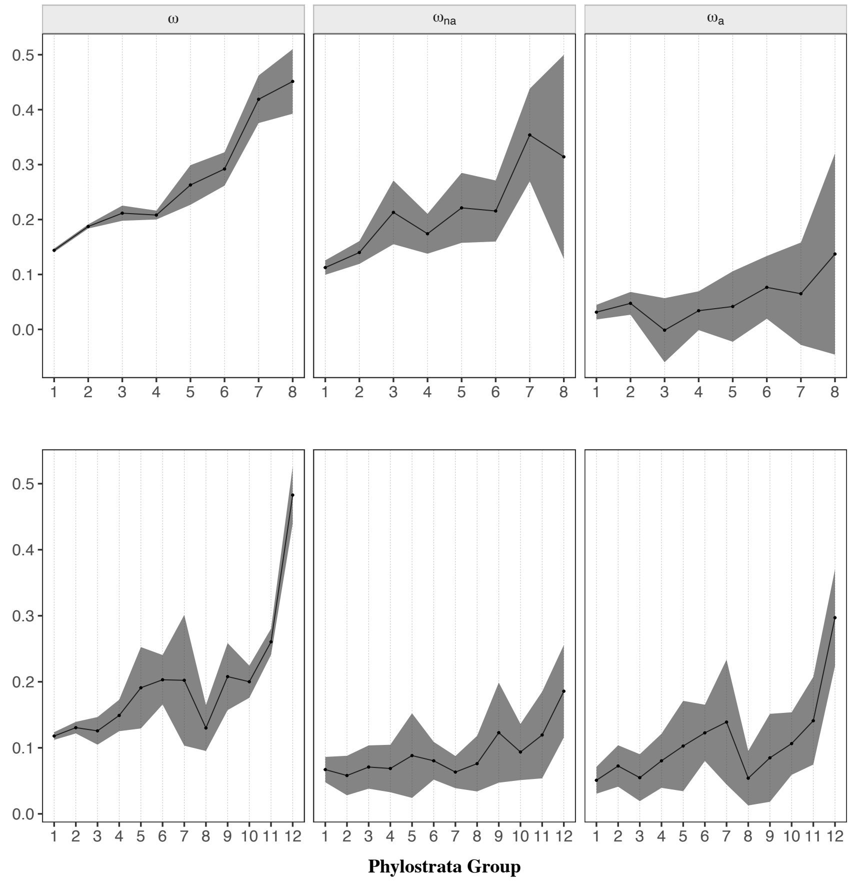


**Figure S4**. Estimates of ω, $\omega_{na}$ and $\omega_a$ plotted as a function of gene age by correcting for the E value on BLAST's searches in *A. thaliana* (top) and *D. melanogaster* (bottom). Mean values of ω, $\omega_{na}$ and $\omega_a$ for each category are represented with the black points. Error bars denote for the 95% confidence interval for each category, computed over 100 bootstrap replicates.

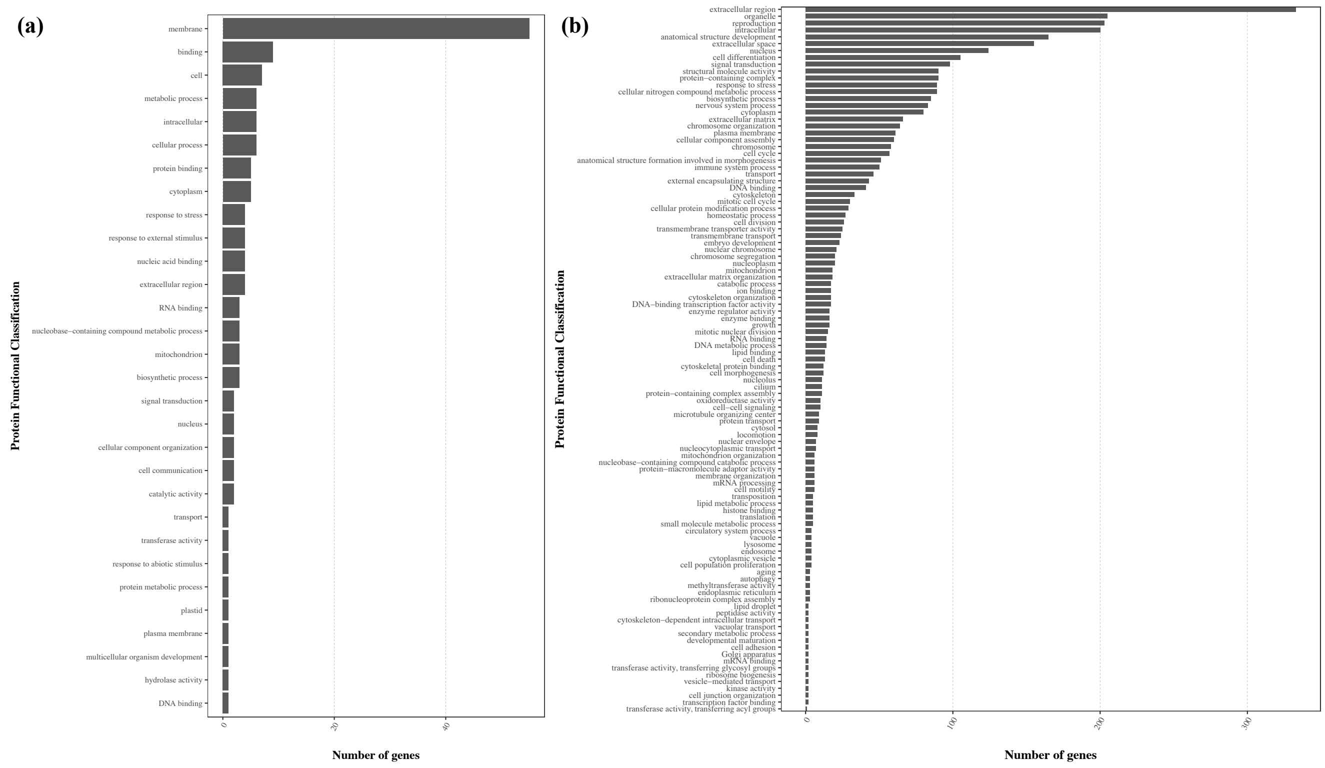


**Figure S5**. Number of young genes for the respective protein function in (a) *D. melanogaster* and (b) *A. thaliana*.

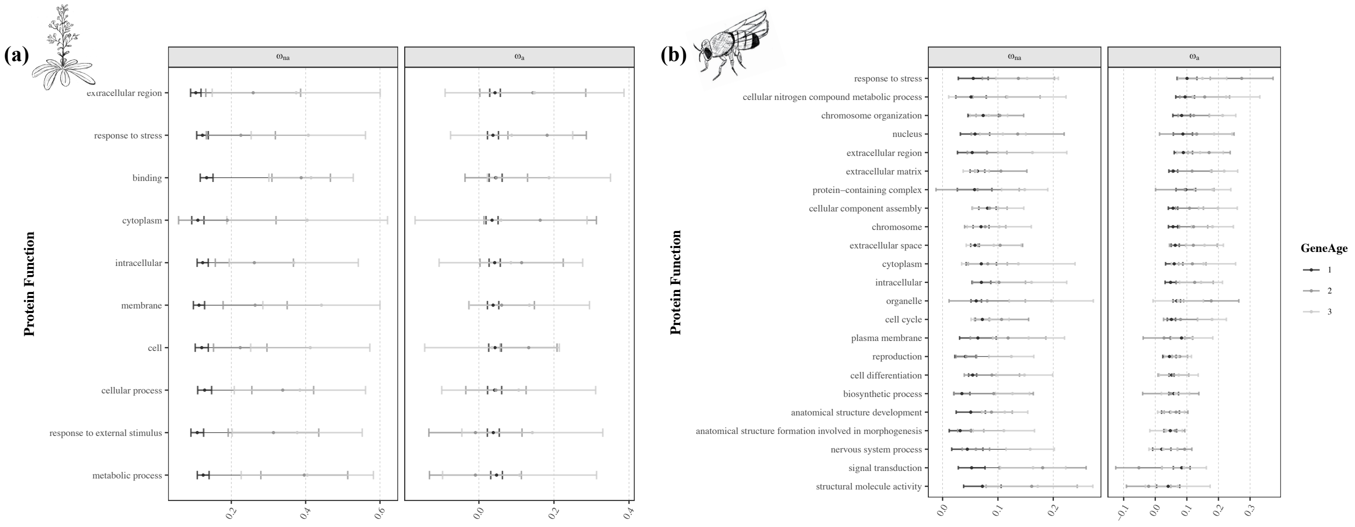


**Figure S6**. Estimates of ω, $\omega_{na}$ and $\omega_a$ plotted as a function of protein function and gene age in (a) A. thaliana and (b) D. melanogaster. Categories are ordered according to the values of $\omega_a$. Gene age categories are ordered from old (1) to young (3). Mean values of ω, $\omega_{na}$ and $\omega_a$ for each class are represented with the black points. Error bars denote the 95% confidence interval for each category, computed over 100 bootstrap replicates.

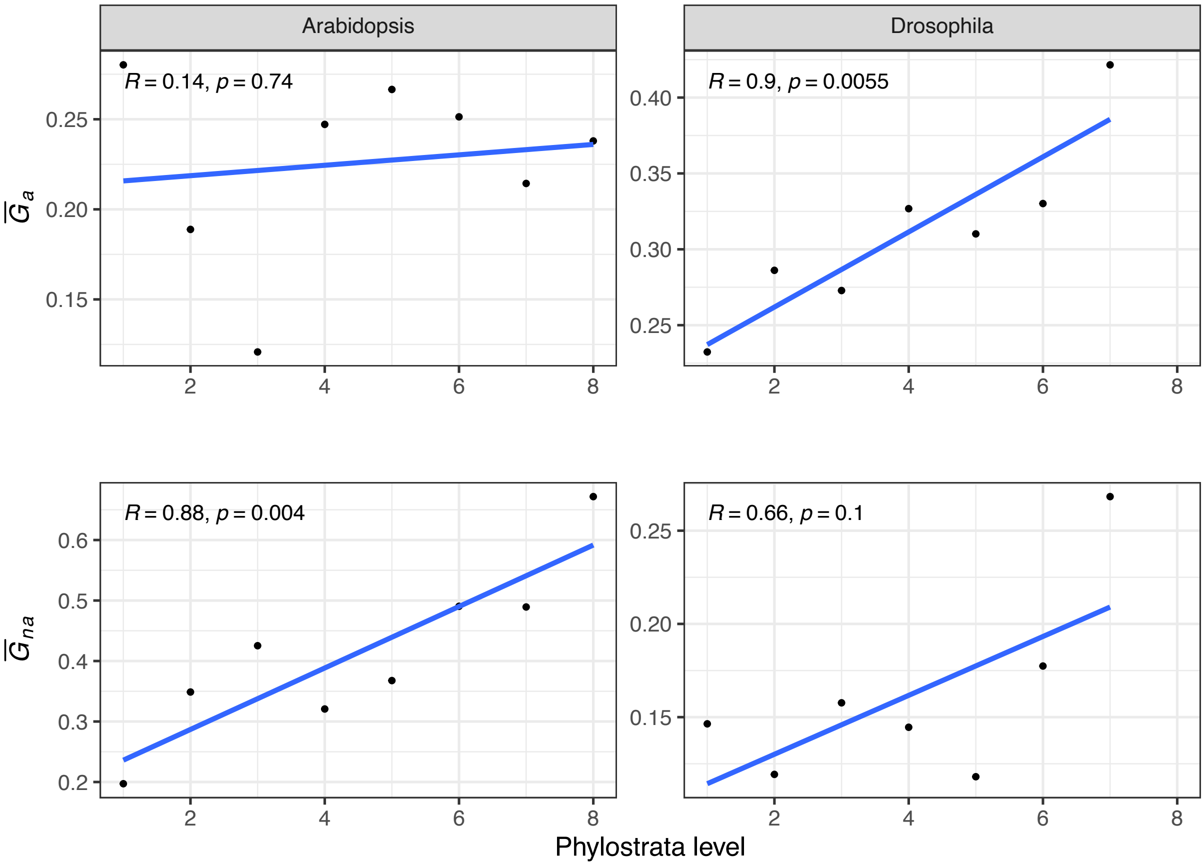


**Figure S7.** Relationship between the proportion of adaptive ($\overline{G_a}$) and non-adaptive ($\overline{G_{na}}$) substitutions and gene age. Each point represents the weighted average for each age category. A linear model was fitted between gene age and Grantham's distances values and is represented with the blue line. Statistical significance was assessed with a Pearson's correlation test and the respective correlation coefficient (R) and p-values (p) are shown in each plot.

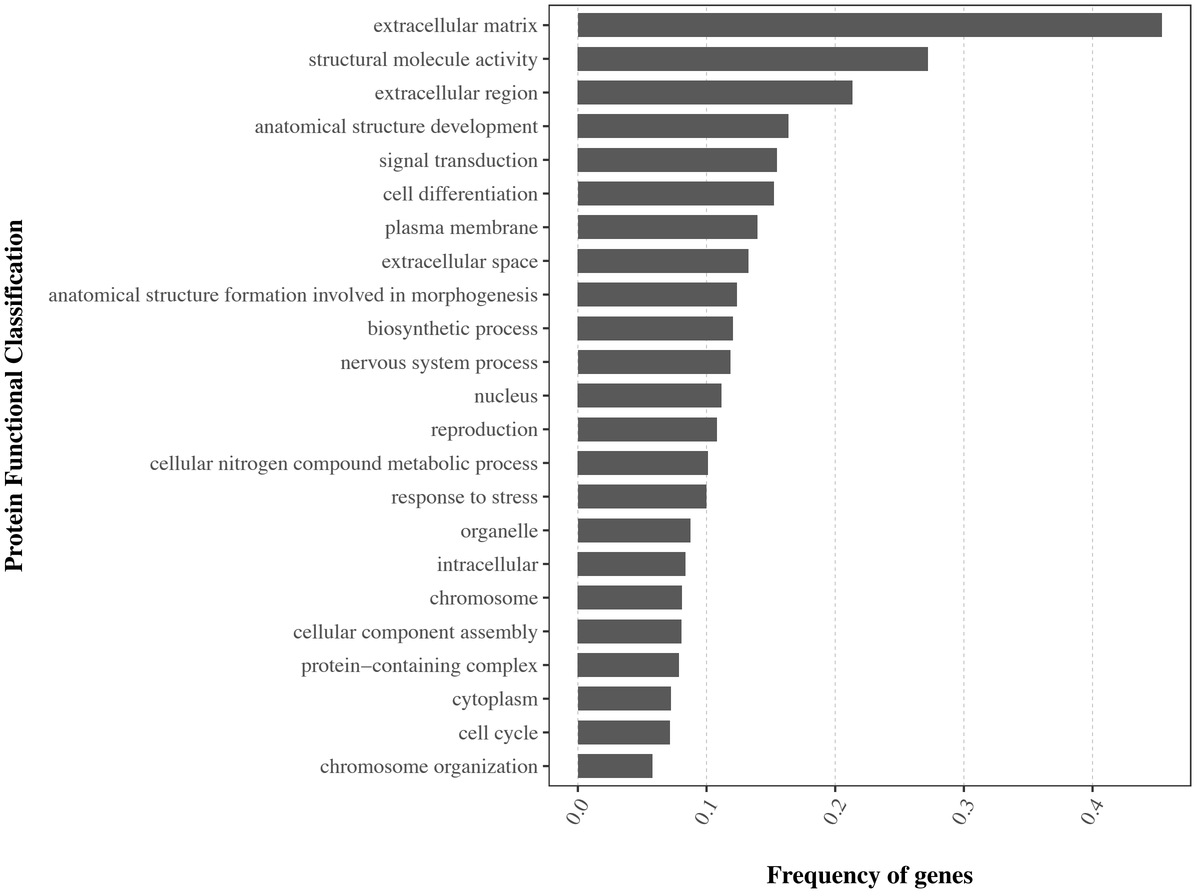


**Figure S8**. Frequency of genes belonging to clades 4-7 for the respective protein function in *D. melanogaster*. The frequency of genes for each functional category was estimated by dividing the number of genes present in these clades by the total number of genes annotated with the respective function.

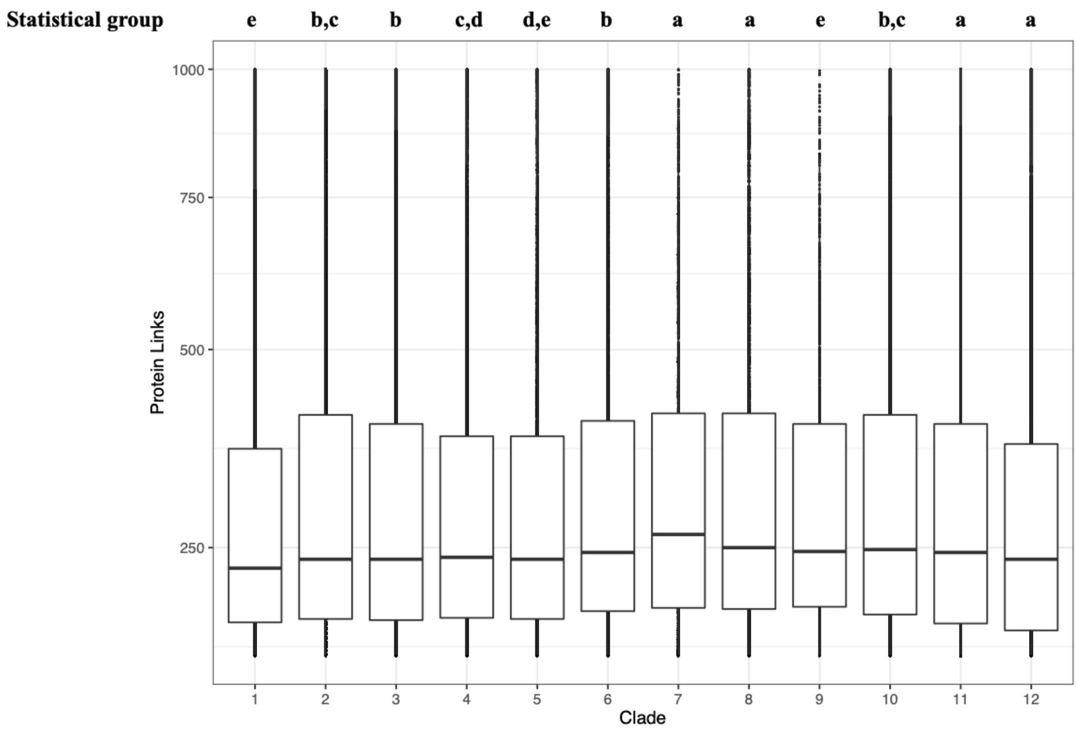


**Figure S9**. Distribution of the degree of PPI values in *D. melanogaster*. The statistical group for each clade is represented. The black line represents the median value of PPI for each clade and black dots denote the outliers of the distribution. The y axis is scaled with a square root function.

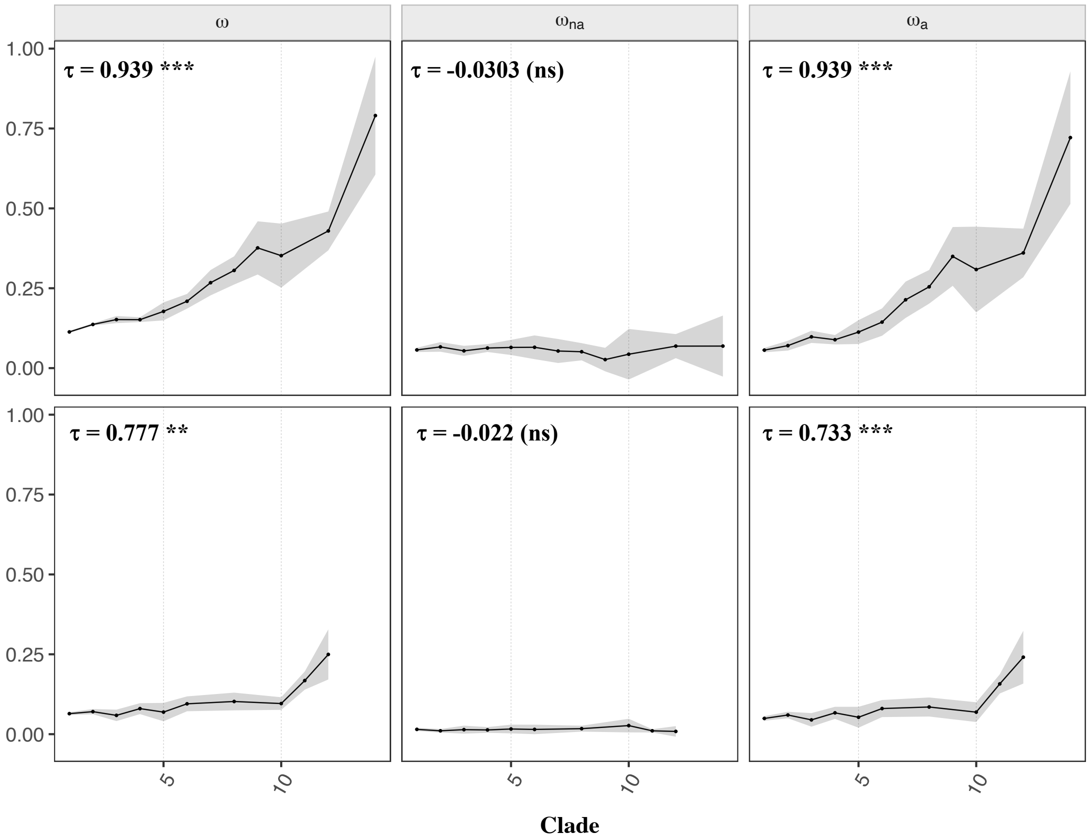


**Figure S10**. Estimates of ω, $\omega_{na}$ and $\omega_a$ plotted as a function of gene age by correcting for the variation in $p_n/p_s$ in *A. thaliana* (top) and *D. melanogaster* (bottom). The Kendall's correlation coefficients are shown with the respective significance (*P < 0.05; **P < 0.01; ***P < 0.001; "." 0.05 ≤ P < 0.10). Legend as in figure S4.
